# RNA-seq analysis of *Triatoma gerstaeckeri* during mite infestation reveals increased metabolic demand and lower reproductive investment

**DOI:** 10.64898/2026.10.06.756886

**Authors:** Alina Cemal, Jacob Bryant, Alekhya Kondragunta, Syeda Farjana Hoque, Kennan Oyen, Sarah A. Hamer, Gabriel L. Hamer, Joshua B. Benoit

## Abstract

Triatomine insects, also known as kissing bugs, serve as vectors for the *Trypanosoma cruzi* parasite, which can cause American trypanosomiasis, also known as Chagas disease. Triatomines are known to be parasitized by mites in natural and colony settings, where mites feed on the arthropod to obtain nutrients and resources for reproduction. This feeding, while necessary for the mite’s survival, likely occurs at the expense of their insect host. We examined the transcriptional response of *Triatoma gerstaeckeri* infested with mites (*Pimeliaphilus* spp.) and an uninfested group using Illumina-based RNA-seq, followed by targeted measurement of nutritional reserves in both groups. Our results show increased expression of factors involved in protein translation, but decreased expression of processes associated with cell development. These results are supported by declines in protein levels in *mite-infested T. gerstaeckeri* and lower expression of reproductive factors, suggesting a distinct reduction in nutrient reserves and reproductive output. These studies provide critical information on the interaction of mites and triatomines while considering the utility of mites as biological control of triatomines in nature or for husbandry protocols to minimize issues with mite infestations of colony individuals.

## Introduction

Members of the subfamily Triatominae (*Reduviidae*), also known as kissing bugs, are a group of insect vectors most notably associated with the transmission of the parasite *Trypanosoma cruzi*, etiologic agent of Chagas disease, a potentially lifelong illness prevalent in Central and South America. Ten million people are estimated to be currently infected with Chagas, and another 30,000 are infected each year (Cousin and Mosser). Nearly 156 species of triatomine insects could pose as vectors for *T. cruzi* (Bern et al. 2019, Guarneri and Lorenzo 2022). *Trypanosoma cruzi* infects a triatomine host during the ingestion of a blood meal from an infected mammalian host. Within the US, triatomines are known to occur in 29 states (Ceccarelli et al. 2018, Curtis-Robles et al. 2018), and populations extend throughout Latin America into Argentina (Beard et al. 2003, Chaves et al. 2023), but are more commonly associated with impoverished and rural communities in Mexico, Central and South America. *Triatoma gerstaeckeri* is a common triatomine species encountered near homes in Texas and northern Mexico and is considered an important vector of *T. cruzi* in this region. Additional vectors of *T. cruzi* in the southern US include *T. rubida* in the southwest (Ribeiro et al. 2012, Martínez-Ibarra et al. 2020) and *T. sanguisuga* (Dumonteil et al. 2020, Dye-Braumuller et al. 2020, Nielsen et al. 2021).

Ectoparasitic mites can impose physiological and fitness costs on their insect hosts (Lehmann 1993, Polak 1996, Luong et al. 2017, Benoit et al. 2020). By feeding on hemolymph or other body fluids, these mites deplete nutrient reserves, impair growth and development, and increase the energetic demands of wound healing and immune responses (Garedew et al. 2004, Benoit et al. 2020). Heavy mite infestations can decrease survival, delay development, reduce reproduction, and alter behaviors such as feeding, dispersal, and mating (Moller 1993, Polak 1996). Ectoparasitic mites may act as vectors of pathogens or facilitate secondary infections by creating wounds that provide entry points for microorganisms (Jaenike et al. 2007, Stone et al. 2024, Skičková et al. 2025). The severity of these impacts depends on factors including mite species, infestation intensity, host life stage, and environmental conditions (Polak and Markow 1995, Rolff et al. 2001, Okabe 2013, Benoit et al. 2026). Consequently, ectoparasitic mites are increasingly recognized as important ecological and evolutionary drivers that influence insect health, population dynamics, and host–parasite interactions.

*Pimeliaphilus* is a genus of mites that are obligate parasites of many arthropods, ranging from scorpions to beetles (Fajfer and Karanth 2021). Species of *Pimeliaphilus* are known to infest triatomine insects in the wild, including *T. gerstaeckeri* (Anderson 1968, Martinez-Sanchez et al. 2007, Dye-Braumuller et al. 2021). Mite infestation in triatomines has been shown to reduce fertility, breeding capacity, and nutritional reserves in triatomine hosts (JosÉ Lino et al. 2004, Martinez-Sanchez et al. 2007). As such, these mites have been suggested for biological control of triatomine vectors, which could help mitigate the burden of Chagas disease (Tian et al. 2024). The presence of mites in triatomine research colonies requires careful management to prevent reduced fecundity or even death following heavy mite infestations (Sutcliffe and Dotson 2024). Despite the negative repercussions of mite infections on the reproductive capacity of triatomine hosts, little is known about how these infestations affect triatomines.

In this study, we examined transcriptional shifts associated with *Pimeliaphilus* sp. mite infestation in *T. gerstaeckeri* to determine how triatomines respond to these ectoparasites. Specifically, we used RNA-seq to determine the molecular mechanisms of *T. gerstaeckeri* infestation with mites. We then assessed protein reserve levels during mite infestation. This study contributes to our growing understanding of how ectoparasites affect their kissing bug host and provides a foundation for future work investigating the possibility of using mites as a biocontrol for triatomine vectors of Chagas disease and improving the management of triatomine colonies.

## Materials and Methods

### *Triatoma gerstaeckeri* collection

*Triatoma gersteackeri* insects were collected in the field in Texas using manual searches, blacklights, and a scent-detection dog as previously described (Curtis-Robles et al. 2015, Christopher et al. 2023). As part of colony maintenance protocols, individual insects collected in the wild are inspected for mites at the time of installation into the colony. Naturally-infested individuals from the same location are housed together in the colony, with apparently uninfested individuals housed separately. Ten mite-infested insects were collected for this study (Fig. 1), including three adult females (F0 generation; Frio/Hidalgo Cos., TX) and seven nymphs (F1 generation; Uvalde Co., TX). We also collected an age- and geography-matched set of ten uninfested females and nymphs. All individuals were euthanized in the freezer.

**Figure 1.**
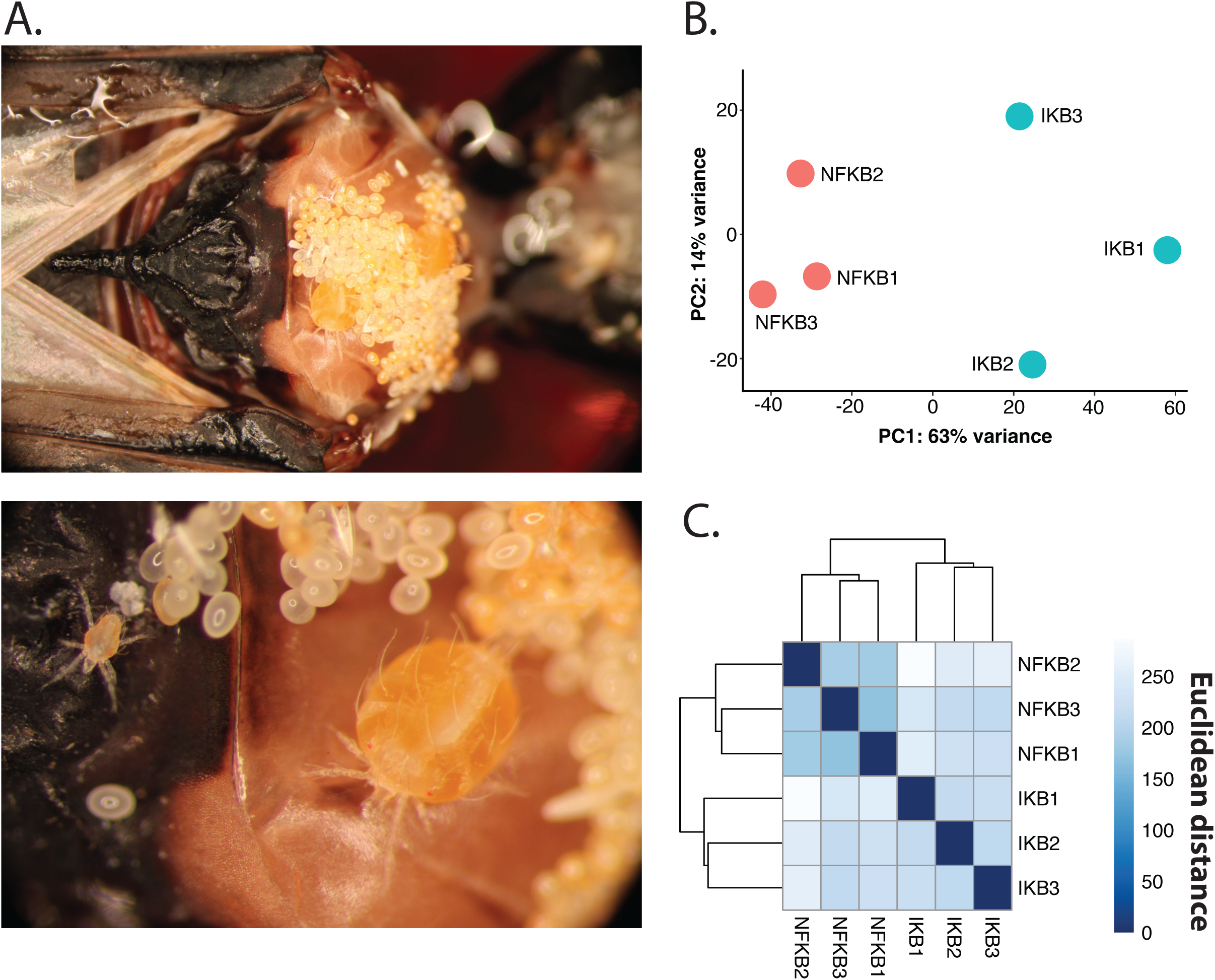
Parasitic mites infesting *Triatoma gerstaeckeri* and transcriptomic analysis of mite infestation. (A) Adult female parasitic mites, *Pimeliaphilus* sp. attached to the thorax of a female *Triatoma gerstaeckeri*. Numerous eggs are deposited in dense, aggregated clusters surrounding the adult mites. The lower panel shows an adult female actively ovipositing, with a larval mite visible to the left, illustrating the rapid and prolific egg-laying behavior of the parasite. (B) Principal component analysis (PCA) of normalized expression profiles. Samples segregate into two distinct clusters along PC1 (63% of the total variance), with PC2 accounting for an additional 14%, indicating differential expression patterns among the analyzed genes. (C) Hierarchical clustering heatmap based on Euclidean distance.

### RNA isolation and sequencing

Females were used to allow for more direct comparison of mite infestation compared to those not harboring mites. RNA was isolated using a TRIzol reagent kit (Invitrogen, Carlsbad, CA) following the manufacturer’s protocol. After isolation, RNA was then treated with DNase I to remove any contaminants, cleaned using the GeneJET microkit (ThermoFisher), and quantified with a Nanodrop (ThermoFisher). Poly(A) libraries were prepared using the TruSeq Stranded mRNA preparation kit at the Cincinnati Children’s Hospital Medical Center. Samples were then barcoded for multiplex analysis, amplified (15-cycle PCR), and libraries sequenced on the Illumina HiSeq 2500 platform.

### De novo assembly for *Triatoma gerstaeckeri*

De novo assembly and RNA-seq studies were conducted based on previously developed methods (Finch et al. 2019, Meibers et al. 2019, Pathak et al. 2022). Reads were quality filtered by trimming low-quality bases (Phred score threshold = 0.05) and removing reads containing ambiguous nucleotides. An additional five nucleotides from the 5′ end and eight nucleotides from the 3′ end were trimmed, and reads shorter than 45 bp were discarded. Read quality before and after filtering was assessed using FastQC, confirming Phred quality scores of approximately 40 across all datasets. Transcriptome assemblies were generated using a combination of CLC Genomics Workbench (Qiagen), Trinity (Grabherr et al. 2011), and Velvet-Oases (Schulz et al. 2012). For all assemblers, the minimum contig length was set to 125 bp, with assembly parameters based on those described (Rosendale et al. 2016). Each assembly was processed individually and in combination for downstream analyses. Protein-coding sequences were predicted using TransDecoder (Haas et al. 2013). Redundant sequences were removed with CD-HIT-EST (Huang et al. 2010) using sequence similarity thresholds of 95%, 93%, and 90%. Assembly completeness was evaluated using BUSCO (Tegenfeldt et al. 2025) with the arthropod reference dataset. A similar process was used for the mites, followed by BLAST to confirm identity, which showed that sequences for ribosomal subunits and cytochrome c oxidase are most closely related to *Pimeliaphilus hemidactyli* (Fajfer and Karanth 2021).

### Differential transcript levels during mite infestation

Samples were analyzed by methods as previously described (Benoit et al. 2025, Tompkin et al. 2026). Raw reads were trimmed for adapters using Trimmomatic (Bolger et al. 2014). Reads shorter than 40 bp were removed from the sample, as were reads with Phred scores smaller than 36. The remaining reads were then assessed for quality with FastQC (Andrews, 2010). Transcript expression was quantified in Sailfish (Patro et al. 2014) and analyzed using DESeq2 (Love et al. 2014) for differential expression. Differentially expressed genes (DEGs) were identified by BLAST in comparison to *Rhodnius prolixus* predicted proteins (Altschul et al., 1990). To assess the putative biological effects of DEGs, functional enrichment analysis was conducted in g:Profiler (Raudvere et al., 2019).

### Assessment of protein content

Protein content was measured in both mite-infested and control experimental groups using previously described methods (Polak et al. 2017, Rosendale et al. 2019). Seven nymphs were selected with and without mites. Mites were removed from the mite-infested experimental group before assay. Samples were then desiccated in a drying oven and then weighed to get the dry weight of each individual. The samples were then homogenized in a 125 µl TET buffer with 1.4-mm ceramic beads in a Bead Blast 24 (Benchmark Scientific). Protein content was then measured in the homogenized samples using a Bradford assay (Thermo Fisher Scientific). Statistical significance was determined using a t-test.

## Results

### *Triatoma gerstaeckeri* de novo assembly

The assembly comprised 93,543 contigs. Assembly completeness was assessed using BUSCO v5.8.0 against the Arthropod_odb10 lineage dataset. The transcriptome recovered 86.7% complete BUSCOs (85.3% single-copy and 1.4% duplicated), with an additional 5.1% fragmented and 8.1% missing BUSCOs, indicating a high level of completeness. Transcriptome annotation was performed by comparing predicted protein sequences from *T. gerstaeckeri* against the *R. prolixus* reference proteome using BLAST; over 52% produced significant matches (E-value < 0.001). All data sets are available through the NCBI Bioproject PRJNA1521935.

### RNA-seq analysis of mite infestation

Comparison of gene expression profiles between *T. gerstaeckeri* infested with ectoparasitic mites and uninfested controls revealed distinct patterns of transcriptional activation and repression associated with mite infestation. Infection was associated with an upregulation of protein synthesis, metabolism, and cellular biosynthesis genes, and reduced expression of DNA replication, cell cycle regulation, signaling, and development genes (Figure 2).

**Figure 2.**
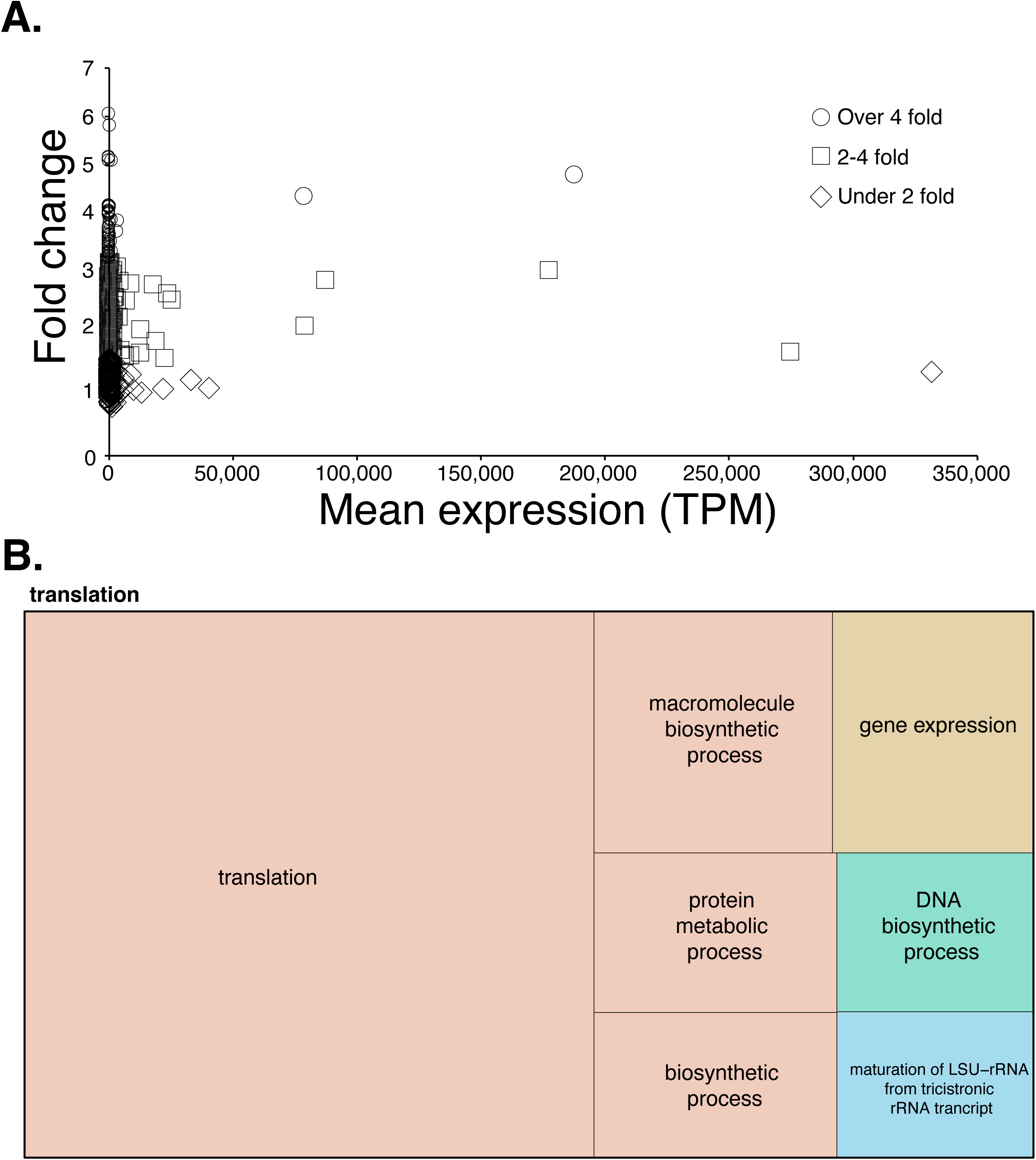
Expression patterns and functional enrichment of genes upregulated during *Pimeliaphilus* sp. infestation of *Triatoma gerstaeckeri.* (A) Scatter plot showing the relationship between mean transcript abundance (transcripts per million, TPM) and fold change of genes upregulated during mite infestation. Each point represents a differentially expressed gene and is categorized by the magnitude of upregulation: over 4-fold (circles), 2–4-fold (squares), and under 2-fold (diamonds). (B) Treemap showing the Gene Ontology (GO) biological process enrichment of genes upregulated during mite infestation. The area of each rectangle is proportional to the relative representation of the corresponding GO term.

Among the upregulated transcripts, genes involved in protein synthesis and biosynthetic processes exhibited the highest transcript abundance and the greatest fold changes (Figure 2). The MA plot (Figure 2A) showed that although most differentially expressed genes were present at relatively low transcript abundance, several highly abundant transcripts (>180,000 TPM) exhibited more than four-fold increases in expression following mite infestation. GO enrichment identified translation as the predominant enriched process, followed by macromolecule biosynthetic process, protein metabolic process, gene expression, DNA biosynthetic process, biosynthetic process, and maturation of LSU-rRNA from tricistronic rRNA transcript (Figure 2B).

The genes contributing to these enriched categories suggest that mite infestation induces a broad increase in protein production and metabolic activity. Among the most strongly upregulated annotated genes were a glutathione peroxidase homolog with thioredoxin peroxidase activity (log₂FC = 6.09), protein takeout-like isoforms (log₂FC = 5.11–5.15), soma ferritin-like (log₂FC = 5.04), a facilitated trehalose transporter Tret1-like (log₂FC = 2.92), UDP-glycosyltransferase UGT5-like (log₂FC = 2.62), and a serine/threonine-protein phosphatase alpha-2 (log₂FC = 2.51) (Table S1). Several highly expressed uncharacterized transcripts also showed strong induction, with expression levels exceeding 180,000 TPM and log₂ fold changes greater than 5.8, indicating that a relatively small number of highly abundant transcripts account for much of the transcriptional response to mite infestation.

In contrast, infestation was associated with widespread downregulation of genes involved in DNA replication and cellular regulatory functions (Figure 3). The MA plot (Figure 3A) showed that most differentially expressed genes had negative fold changes, with several transcripts decreasing by up to 8-fold. GO enrichment indicated that these downregulated genes were significantly enriched for DNA metabolic processes, RNA-templated DNA biosynthesis, protein phosphorylation, aromatic amino acid biosynthesis, signal transduction, cellular response to stimulus, regulatory ncRNA-mediated gene silencing, microtubule-based movement, cell communication, cell cycle, lipid catabolic process, and response to stimulus (Figure 3B).

**Figure 3.**
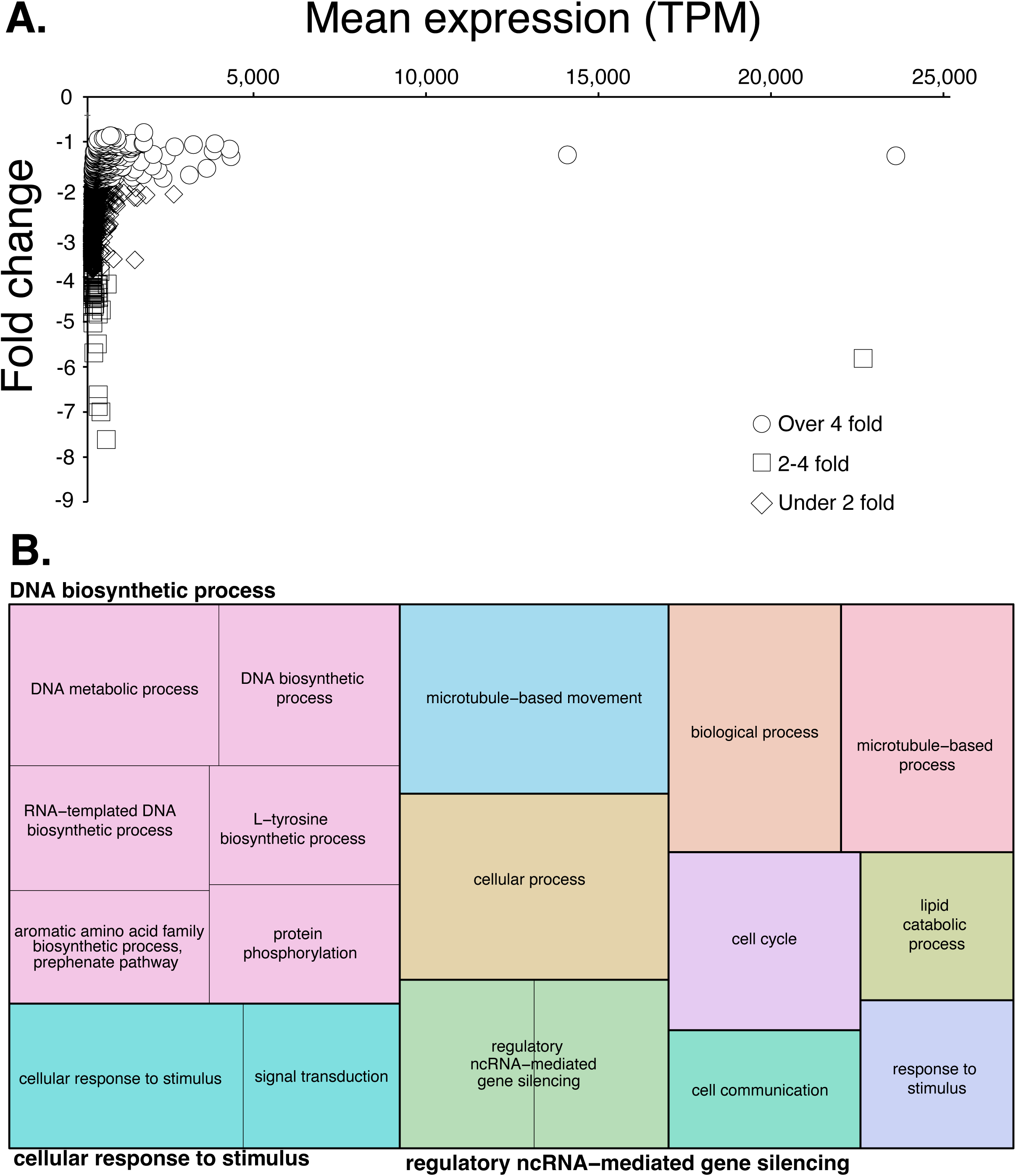
Expression patterns and functional enrichment of genes downregulated during *Pimeliaphilus* sp. infestation of *Triatoma gerstaeckeri*. (A) Scatter plot showing the relationship between mean transcript abundance (transcripts per million, TPM) and fold change of genes downregulated during mite infestation. Each point represents a differentially expressed gene and is categorized by the magnitude of downregulation: over 4-fold (circles), 2–4-fold (squares), and under 2-fold (diamonds).. (B) Treemap showing the Gene Ontology (GO) biological process enrichment of genes downregulated during mite infestation. The area of each rectangle is proportional to the relative representation of the corresponding GO term.

Downregulation of several genes involved in cell proliferation, development, genome maintenance, and transcriptional regulation was observed (Table S2). Transcripts encoding an ecdysoneless cell cycle regulator (log₂FC = –1.21), ecdysone-induced protein 78C (log₂FC = –3.44), and the tudor domain-containing protein qin (log₂FC = –1.21) were significantly reduced in mite-infested insects. We also observed decreases in genes involved in RNA silencing and genome maintenance, including the Piwi-like protein Siwi, the 5′–3′ exoribonuclease pacman, the mutagen-sensitive 101, and the SET domain-containing protein upSET. Genes associated with metabolism and signaling, including phenylalanine hydroxylase, anoctamin-7-like, and the trypsin-like serine protease masquerade, were also downregulated. There was a notable reduction in vitellogenin protein levels (Table S2). Several of the largest decreases (log₂FC = –5.2 to –7.8) occurred in transcripts lacking functional annotation, suggesting that additional, uncharacterized genes may contribute to the host response to ectoparasitic mite infestation.

Overall, *T. gerstaeckeri* infestation by parasitic mites was associated with a marked shift in gene expression pathways involving translation, protein synthesis, antioxidant defense, and metabolism, then decreased expression of genes involved in DNA replication, cell cycle progression, intracellular signaling, and gene regulatory processes. These findings show that mite infestation substantially reprograms the triatomine transcriptome, consistent with a physiological response involving enhanced biosynthetic activity coupled with suppression of cellular growth and proliferation.

### Protein levels during mite infestation

Associations between mite parasitism and host nutrient reserves demonstrated that protein content was significantly greater in the control group (ca. 1142.75 +/- 77 µg protein/mg dry mass) than in the mite-infested experimental group (ca. 845.87 +/- 51 µg protein/mg dry mass), representing an average reduction of 296.88 µg protein/mg dry mass (ANOVA, P < 0.05). This substantial decrease in protein reserves suggests that mite parasitism imposes a considerable energetic burden on the host.

## Discussion

This study investigated the effects of ectoparasitic mite infestation on the physiological condition and transcriptional responses of the triatomine *T. gerstaeckeri*. Mite infestation was associated with reduced host protein reserves while inducing widespread changes in gene expression, indicating that parasitized insects undergo substantial metabolic reorganization. Transcriptome analysis showed that infested insects exhibited increased expression of genes involved in translation, antioxidant defense, and cellular biosynthesis. In contrast, genes associated with DNA replication, cell cycle progression, intracellular transport, and gene regulation were broadly downregulated. Together, these findings suggest that mite infestation imposes an energetic burden that alters host resource allocation and physiological function.

One of the most striking transcriptional responses to mite infestation was the upregulation of genes involved in translation. GO enrichment identified translation as the main biological process, accompanied by increased expression of genes involved in macromolecule biosynthesis, protein metabolism, and gene expression. Several of the most strongly induced genes encoded glutathione peroxidase, ferritin, trehalose transporters, glycosyltransferases, and protein phosphatases, all of which are associated with oxidative stress protection, nutrient metabolism, and cellular homeostasis (Chaitanya et al. 2016, Tellis et al. 2023). Increased expression of glutathione peroxidase and ferritin likely reflects enhanced protection against oxidative damage resulting from tissue injury caused by mite feeding (Sterkel et al. 2017). Importantly, blood ingestion yields substantial oxidative stress, which means this stress is exacerbated following feeding. Of interest is that these factors suggest that blood ingestion or starvation are likely to be altered, but that was not observed in the *Pimeliaphilus plumifer-Meccus pallidipennis* system (Martinez-Sanchez et al. 2007). These differences could reflect differences between organismal systems, such that responses differ depending on triatomine-mite systemic differences, warranting further investigations.

Mite-infested *T. gerstaeckeri* exhibited significantly lower total protein reserves than uninfested controls. Protein synthesis is energetically costly and is required for tissue repair, cellular maintenance, and production of proteins necessary for recovery following parasite-induced damage (Rivero and Ferguson 2003, Mortimer and Schlenke 2025). *Bombyx mori* infected with parasites deplete nutrient reserves while increasing amino acid metabolism and biosynthetic activity to sustain physiological function (Dai et al. 2022). Likewise, *Drosophila* infected with ectoparasitic mites have increased metabolic activity to support energetically expensive defense mechanisms and tissue repair (Luong et al. 2017, Benoit et al. 2020, Brophy and Luong 2021). The elevated translation and biosynthesis observed in *T. gerstaeckeri* likely reflect an attempt to replace proteins and maintain cellular function despite continual nutrient loss from mite feeding. Consistent with this interpretation, mite infestation was associated with approximately 26% lower total protein reserves and similar observations of lower nutrient reserves have been observed in other insect-mite systems (Yang and Cox-Foster 2007, Benoit et al. 2020)

DNA replication, cell cycle regulation, intracellular signaling, and cellular transport were broadly downregulated in infested insects. Collectively, these changes suggest that infested insects reduce investment in cellular proliferation and developmental processes while prioritizing immediate physiological maintenance (Schwenke et al. 2016, Dolezal et al. 2019). The observed reduction in microtubule-based processes may have important implications for cellular function during infestation. Microtubules form the structural framework that supports the intracellular trafficking of proteins, vesicles, organelles, and signaling molecules and are essential for maintaining cellular organization (Burute and Kapitein 2019). Reduced expression of genes associated with microtubule organization and intracellular transport suggests diminished cellular activity and may reflect an energy-conserving strategy during chronic parasitic stress. Alternatively, repeated feeding damage inflicted by mites could disrupt tissue integrity and cytoskeletal maintenance, indirectly impairing intracellular transport. Similar alterations in cytoskeletal organization have been reported in other host-pathogen interactions (Gruenheid and Finlay 2003), although functional consequences vary by parasite and host system. Additional studies examining tissue-specific responses and cytoskeletal architecture are needed to determine the mechanisms underlying these transcriptional changes.

Expression of vitellogenin was reduced in mite-infested *T. gerstaeckeri*, suggesting that mite infestation disrupts host nutrient metabolism and resource allocation. Most likely, depleted protein reserves may limit the host’s ability to maintain reproduction. A limitation of our study is that we examined nymphs, not adult females, for protein content, but similar reductions in protein levels may be expected in females. A reduction in egg production was observed in the *P. plumifer*-*M. pallidipennis* system (Martinez-Sanchez et al. 2007), where fewer viable eggs were laid by females infected with mites. This is common, as ectoparasitic mites reduce insect reproduction because the host must pay a nutritional cost to respond to mite presence and feeding (Benoit et al. 2020, Scaramella et al. 2023). Importantly, many insect populations could develop resistance to mites, but resistance mechanisms often come at a cost (Polak et al. 2023, Benoit et al. 2025). A survey of 403 wild-caught triatomines from Arizona and New Mexico found that 13.2% were infested by mites (*T. rubida* and *T. protracta*) (Dye-Braumuller et al. 2021), suggesting that tolerance rather than resistance may allow higher reproduction.

Overall, the transcriptomic response suggests that mite infestation induces a broad physiological shift rather than activation of a single stress pathway. The concurrent depletion of host protein reserves and reduced vitellogenin expression further demonstrate that this response occurs despite declining nutritional conditions. These results support that ectoparasitic mites impose a substantial energetic cost on triatomines, leading to altered resource allocation that favors short-term maintenance and tissue repair over reproduction. Future studies should determine whether these transcriptional responses are localized to tissues directly damaged by mite feeding or represent systemic physiological changes throughout the insect. Investigating tissue-specific gene expression, immune function, and metabolic pathways during infestation will provide further insight into how ectoparasitic mites influence the biology of *T. gerstaeckeri* and other triatomine vectors.

## Acknowledgements

Illumina sequencing was performed at Cincinnati Children’s Hospital Medical Center DNA Core. Partial funding for computational resources was provided by the National Institute of Allergy and Infectious Diseases of the National Institutes of Health (R01AI148551 to J.B.B.) and the National Science Foundation (NSF, DEB1654417 to J. B.B.). Triatomine colony maintenance and mite detection were conducted by Keswick Killets, Jillian Wormington, Justin Bejcek, Rachel Curtis and Alyssa Meyers, with funding from the Texas A&M AgriLife Research and the Texas EcoLab Program. The funding agencies had no role in the study design, data collection, and interpretation, or the decision to submit the work for publication. Opinions contained in this publication do not reflect the opinions of the funding agencies. We declare no competing financial interests.

---

**Table S1** - Genes with enriched transcript levels during *Pimeliaphilus* mite infestation of *Triatoma gerstaeckeri*. Significance determined with DESeq2 (FDR, P < 0.05).

**Table S2** - Genes with lower transcript levels during *Pimeliaphilus* mite infestation of *Triatoma gerstaeckeri*. Significance determined with DESeq2 (FDR, P < 0.05).

